# Fungal microbiomes are determined by host phylogeny and exhibit widespread associations with the bacterial microbiome

**DOI:** 10.1101/2020.07.07.177535

**Authors:** Xavier A. Harrison, Allan D. McDevitt, Jenny C. Dunn, Sarah Griffiths, Chiara Benvenuto, Richard Birtles, Jean P. Boubli, Kevin Bown, Calum Bridson, Darren Brooks, Samuel S. Browett, Ruth F. Carden, Julian Chantrey, Friederike Clever, Ilaria Coscia, Katie L. Edwards, Natalie Ferry, Ian Goodhead, Andrew Highlands, Jane Hopper, Joseph Jackson, Robert Jehle, Mariane da Cruz Kaizer, Tony King, Jessica M. D. Lea, Jessica L. Lenka, Alexandra McCubbin, Jack McKenzie, Bárbara Lins Caldas de Moraes, Denise B. O’Meara, Poppy Pescod, Richard F. Preziosi, Jennifer K. Rowntree, Susanne Shultz, Matthew J. Silk, Jennifer E. Stockdale, William O. C. Symondson, Mariana Villalba de la Pena, Susan L. Walker, Michael D. Wood, Rachael E. Antwis

## Abstract

Interactions between hosts and their resident microbial communities are a fundamental component of fitness for both agents. Though recent research has highlighted the importance of interactions between animals and their bacterial communities, comparative evidence for fungi is lacking, especially in natural populations. Using data from 49 species, we present novel evidence of strong covariation between fungal and bacterial communities across the host phylogeny, indicative of recruitment by hosts for specific suites of microbes. Using co-occurrence networks, we demonstrate that fungi form critical components of putative microbial interaction networks, where the strength and frequency of interactions varies with host taxonomy. Host phylogeny drives differences in overall richness of bacterial and fungal communities, but the effect of diet on richness was only evident in mammals and for the bacterial microbiome. Collectively these data indicate fungal microbiomes may play a key role in host fitness and suggest an urgent need to study multiple agents of the animal microbiome to accurately determine the strength and ecological significance of host-microbe interactions.

**SIGNIFICANCE STATEMENT:** Microbes perform vital metabolic functions that shape the physiology of their hosts. However, almost all research to date in wild animals has focused exclusively on the bacterial microbiota, to the exclusion of other microbial groups. Although likely to be critical components of the host microbiome, we have limited knowledge of the drivers of fungal composition across host species. Here we show that fungal community composition is determined by host species identity and phylogeny, and that fungi form extensive interaction networks with bacteria in the microbiome of a diverse range of animal species. This highlights the importance of microbial interactions as mediators of microbiome-health relationships in the wild.

## INTRODUCTION

Multicellular organisms support diverse microbial communities critical for physiological functioning, immunity, development, evolution and behaviour (1–3). Variability in host-associated microbiome composition may explain asymmetries among hosts in key traits including susceptibility to disease (4, 5), fecundity (6), and resilience to environmental change (7). Although the microbiota is a complex assemblage of bacteria, fungi, archaea, viruses and protozoa, the overwhelming majority of research has focused solely on the bacterial component (8, 9). Although relatively well documented in soils and plants (10–13), relatively few studies have examined the dynamics of non-bacterial components of the microbiome in animal hosts (but see (14–16)), especially in non-model or wild systems. As such, our current understanding of host-microbe interactions is skewed by a bacteria-centric view of the microbiome. Although not well understood, there is growing evidence that the fungal microbiota, termed the ‘mycobiome’, may drive diverse functions such as fat, carbon and nitrogen metabolism (17, 18), degradation of cellulose and other carbohydrates (19), pathogen resistance (20), initiation of immune pathways and regulation of inflammatory responses (9, 21), and even host dispersal (22).

Host phylogeny has repeatedly been shown to be an important predictor of bacterial microbiome structure in multiple vertebrate clades, a phenomenon known as ‘phylosymbiosis’ (23–27). This phenomenon often reflects phylogenetic patterns in life history traits, such as diet, physiology or spatial distribution (23–27). However, evidence of phylosymbiosis, and its drivers, in other microbial kingdoms or domains is lacking. Addressing this major gap in our knowledge is crucial as we likely underestimate the strength and importance of coevolution between animal hosts and their resident communities, particularly in the context of cross-kingdom interactions within the microbiome (28).

Here we used ITS and 16S rRNA gene amplicon sequencing to characterise fungal and bacterial communities of primarily gut and faecal samples from 49 host species across eight classes, including both vertebrates and invertebrates (Table S1). We predicted that both fungal and bacterial microbiomes demonstrated strong signals of phylosymbiosis across the broad host taxonomic range tested. Specifically, we predicted that patterns of phylosymbiosis within microbial kingdoms will also drive significant positive covariance in patterns of microbial community structure between microbial kingdoms within individual hosts, suggestive of evolutionary constraints that favour co-selection of specific bacterial and fungal communities in tandem. We also used network analysis to identify key bacteria-fungi interactions whilst quantifying variation in the frequency and strength of bacteria-fungi interaction networks across host taxonomic groups. Finally, we tested the prediction that cross-kingdom phylosymbiosis may be partially driven by similarity in host dietary niche across the 32 bird and mammal species sampled.

## RESULTS

### Fungal and Bacterial Microbiome Diversity Varies with Host Phylogeny

Our data revealed consistent patterns in fungal and bacterial alpha diversity across host taxonomic groups. Bacterial community alpha-diversity was generally greater than, or similar to, fungal community alpha-diversity at the host species level (Fig. 1A), although two species exhibited greater fungal diversity than bacterial (great tit, tsetse fly; Fig. 1A). Comparisons between microbial richness values within individuals (i.e., *relative* richness) using a binomial GLMM supported these patterns, indicating that bacterial richness was higher on average than fungal in 80% of individuals [95% credible interval (CI) 0.55 - 0.95]. When conditioning on Class, samples from both Mammalia and Insecta were more likely to have higher bacterial diversity than fungal diversity (credible intervals not crossing zero on the link scale). Mammalia were more likely to have higher bacterial relative to fungal diversity than Aves in our study organisms (mean difference in probability 22.9% [1.6 - 45.7%]).

**FIGURE 1.**
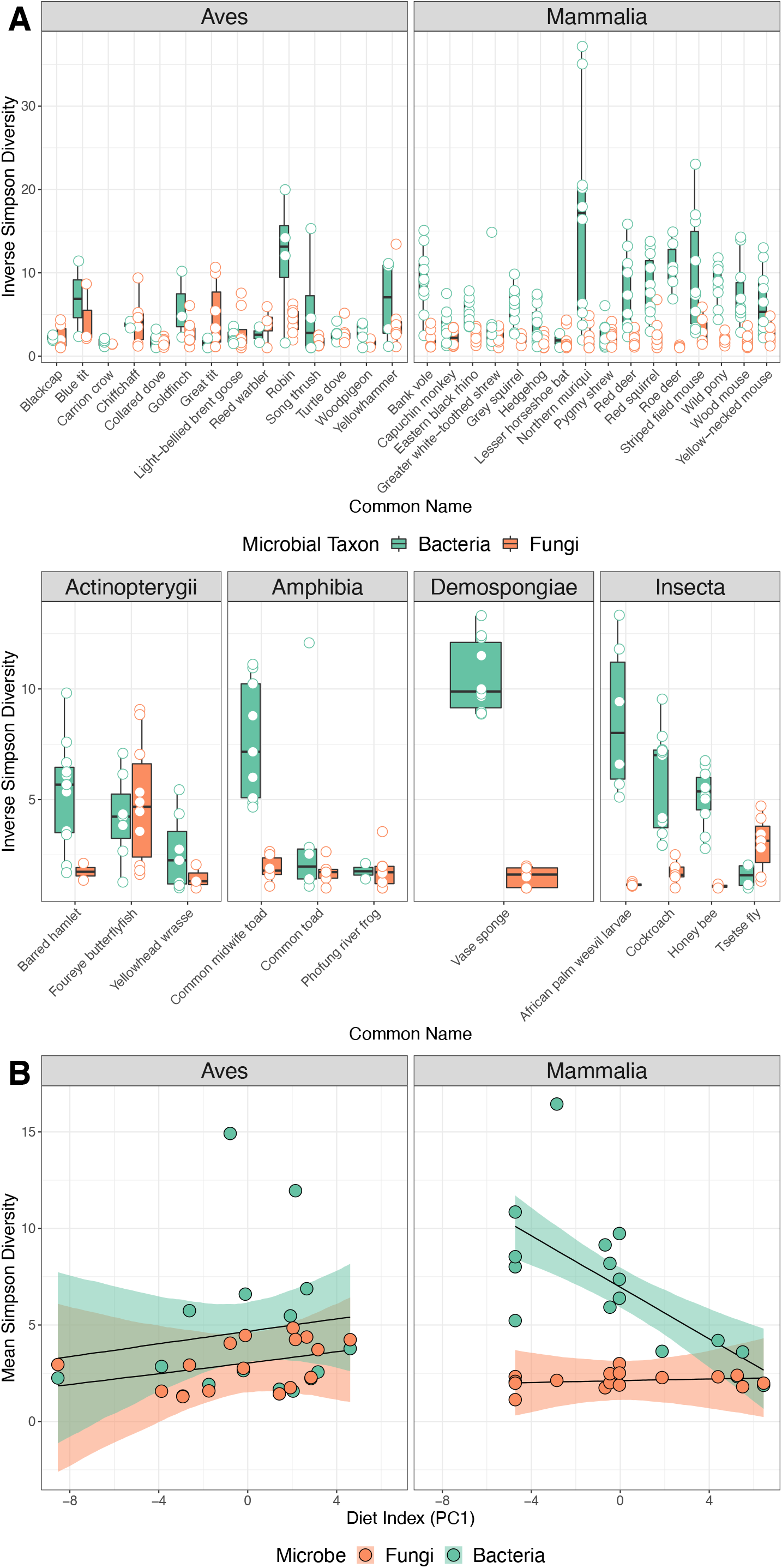
Host phylogeny and diet as predictors of host bacterial and fungal alpha diversity. **(A)** Boxplots and raw data (points) of inverse Simpson indices for bacterial (green) and fungal (orange) communities across a range of host species. **(B)** Raw data (points) and model predictions (shaded area and lines) of models examining the relationship between host diet and microbiome alpha diversity. In mammals, an increase the in the amount of plant material in the diet (more negative PC1 values) drives increases in richness. There was no corresponding relationship between diet and richness for fungi in mammals, nor for bacteria and fungi in birds. Shaded areas represent 95% credible intervals.

Variation among species in this model explained 19.5% [7.3 - 31.2%] of the variation in relative microbial richness. Using a bivariate model with both fungal and bacterial diversity as response variables to examine patterns of absolute microbial richness across host taxonomy, only Mammalia exhibited bacterial diversity that was consistently higher than fungal diversity when controlling for variation among species (mean difference in index 5.16; [3.33 - 6.96]). There was no evidence of positive covariance between fungal and bacterial richness values at the species level (mean correlation 0.3, 95% credible intervals −0.55 - 0.86), suggesting that high diversity of one microbial group does not necessarily reflect high diversity of the other. The bivariate model also revealed that species identity explained 33.9% [22.2 – 44.2%] of variation in bacterial diversity, and 22.4% [9.8 – 35.5%] of variation in fungal diversity.

Phylogenetic analyses supported these general patterns (Fig. S2). For fungi, we detected phylogenetic signal in patterns of both Inverse Simpson index (C_mean_ = 0.22, p = 0.021) and number of observed amplicon sequence variants (ASVs) (C_mean_ = 0.26, p = 0.016). For bacteria, phylogenetic signal was evident for number of ASVs (C_mean_ = 0.28, p = 0.016) but not inverse Simpson index (C_mean_ = 0.114, p = 0.100).

### Limited Evidence of Covariation Between Host Diet and Fungal Microbiome

#### Alpha Diversity

Models exploring the influence of diet on microbial richness yielded mixed results. In mammals, only a relationship between *bacterial* richness and diet was evident (interaction between microbe (fungi vs bacteria) and the primary axis of a PCA of dietary variation; Fig 1B). This indicates that bacterial alpha diversity increases in tandem with the proportion of plant matter in the diet. However, this relationship was absent in birds (Fig. 1B). Similarly, there was no relationship between *fungal* richness and diet for birds or mammals (credible intervals for slopes all include zero).

#### Beta Diversity

Patterns of variation in microbial community *structure* broadly followed those for alpha diversity above. While for mammals there was a significant correlation between host-associated bacterial community composition and diet (r = 0.334, p = 0.002), and a near-significant relationship between fungal community composition and diet (r = 0.142, p = 0.067), for birds there was no significant relationship between dietary data and bacterial community composition (r = 0.087, p = 0.211) or fungal community composition (r = 0.026, p = 0.386). Further, taxonomic differences in microbiome composition based on differences in crude dietary patterns were not clear for either bacteria or fungi when the microbiome composition was visualised at the family level (Figs. S3, S4). That said, Alphaproteobacteria and Eurotiomycete fungi were notably absent from species that primarily ate vegetation (i.e. grasses etc) and Neocallimastigomycete fungi were the predominant fungal class associated with two out of four of these host species (Figs. S3, S4).

### Strong Evidence of Correlated Phylosymbiosis in Both Microbial Groups

Our data revealed consistent variation in fungal and bacterial community structure across the host phylogeny (Fig. 2A). PERMANOVA analyses on centred-log ratio (CLR) transformed ASV abundances revealed significant phylogenetic effects of host class, order and species, as well as effects of sample storage and library preparation protocol for both microbial groups (Table 2; Figs. S5 & S6). For both bacteria and fungi, host species identity explained more variation than host class or order, and this pattern remained when re-running the models without sample preparation protocol effects, though this inflated the estimate of R^2^ for all taxonomic groupings (Table 2).

**FIGURE 2.**
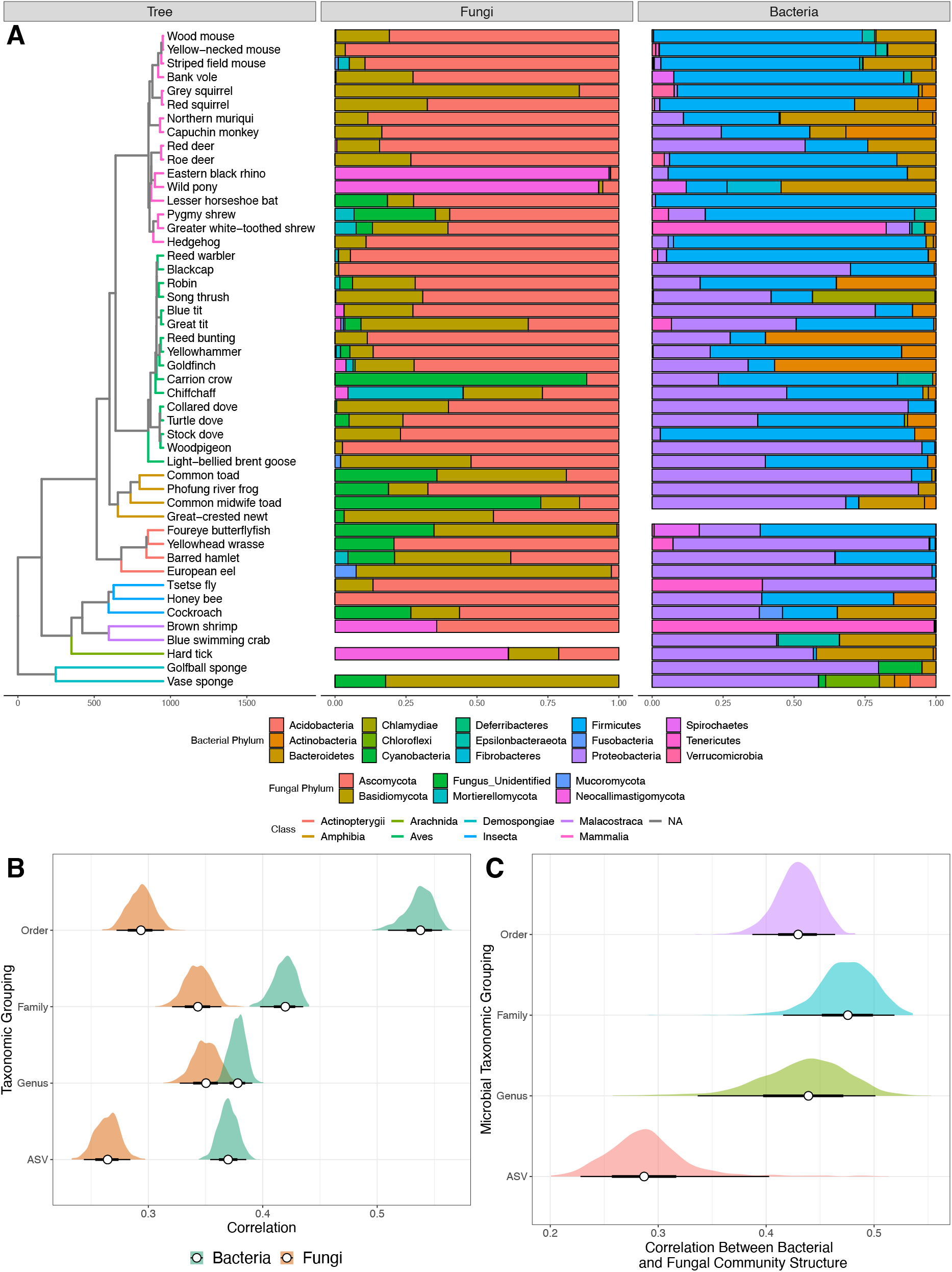
**(A)** Phylogenetic tree of host species, with branches coloured by class and node points coloured by order. Barplots show proportional composition of fungal and bacterial phyla for each host species, aligned to tree tips. **(B)** Correlation between microbial and host genetic distances (generated from the phylogenetic tree in A) for both bacteria (green) and fungi (orange) across all host species. Microbial taxonomy was either raw ASVs or grouped into higher taxonomic levels. Aggregation to higher taxonomy tended to result in higher correlations for both microbial groups, and the correlation was always stronger in bacteria. **(C)** Correlation between fungal and bacterial community structure derived from Procrustes rotation on PCA ordinations of each microbial group. Microbial communities were aggregated at various taxonomic groupings (order, family, genus), or as raw Amplicon Sequence Variant (ASV) taxonomy. For both B and C, distributions of correlation values were generated using resampling of 90% of available samples for that microbial group to generate 95% intervals (shaded areas on graphs). Empty bars in panel 2A mean samples were not available for a particular species and so would not have been included in the calculations in panel B or C.

**TABLE 2.** PERMANOVA results for (a) fungi and (b) bacteria of factors explaining variation in microbial community structure. Terms were added in the order shown in the table to marginalise effects of sample storage and preparation protocols before calculating % variance explained for taxonomic groupings. Species ID was the dominant source of variation in the data for both taxonomic groups, but there were also strong effects of sample storage and wet lab protocol, particularly for bacteria.

| (a) FUNGI |  |  |  | Taxonomic Effects Only |  |  |
| --- | --- | --- | --- | --- | --- | --- |
| Predictor | df | R<sup>2</sup> | p value | df | R<sup>2</sup> | p value |
| Sample Type | 7 | 0.05 | 0.001 |  |  |  |
| Tissue Storage | 5 | 0.04 | 0.001 |  |  |  |
| Extraction Kit | 7 | 0.07 | 0.001 |  |  |  |
| Class | 2 | 0.02 | 0.001 | 6 | 0.05 | 0.001 |
| Order | 6 | 0.05 | 0.001 | 13 | 0.12 | 0.001 |
| Species | 18 | 0.09 | 0.001 | 26 | 0.14 | 0.001 |
| Residuals | 303 | 0.68 |  | 303 | 0.68 |  |

| (b) BACTERIA |  |  |  | Taxonomic Effects Only |  |  |
| --- | --- | --- | --- | --- | --- | --- |
| Predictor | df | R<sup>2</sup> | p value | df | R<sup>2</sup> | p value |
| Sample Type | 6 | 0.06 | 0.001 |  |  |  |
| Tissue Storage | 6 | 0.16 | 0.001 |  |  |  |
| Extraction Kit | 7 | 0.12 | 0.001 |  |  |  |
| Class | 2 | 0.02 | 0.001 | 6 | 0.09 | 0.001 |
| Order | 6 | 0.09 | 0.001 | 12 | 0.21 | 0.001 |
| Species | 18 | 0.12 | 0.001 | 27 | 0.27 | 0.001 |
| Residuals | 273 | 0.42 |  | 273 | 0.42 |  |

**TABLE 2:**
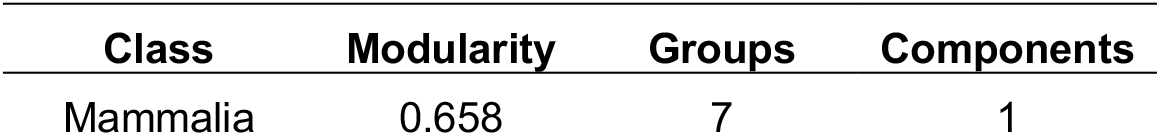
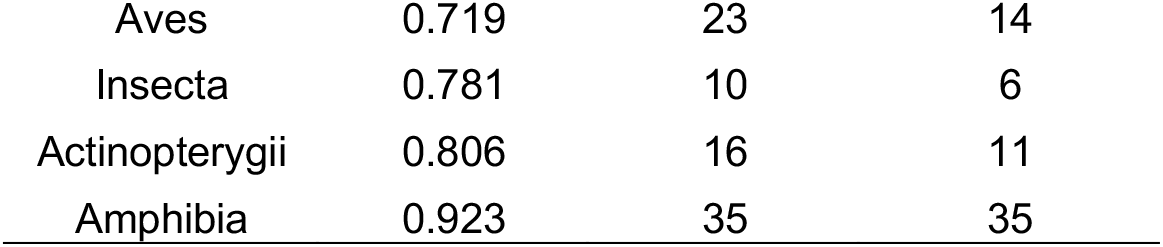
Network statistics from class-specific microbial networks in Figure 3 in the main manuscript. ‘Modularity’ and ‘Groups’ statistics are derived from the cluster_fast_greedy function applied to *igraph* network objects. ‘Components’ data were extracted directly from the networks. Modularity was positively correlated with both number of groups (cor = 0.76) and number of components (cor = 0.86).

Consistent with our predictions, the similarity between the microbial communities of a given pair of host species was proportional to the phylogenetic distance between them (e.g. ASV level: fungal cor. = 0.26; p = 0.001; bacterial cor. = 0.37; p = 0.001; Fig. 2B). Correlations for both bacterial and fungal communities became stronger when aggregating microbial taxonomy to broader taxonomic levels (Fig. 2B). Notably, the bacterial correlation was stronger than the fungal equivalent at most taxonomic levels (Fig. 2B), indicating stronger patterns of phylosymbiosis for bacteria.

We also detected a strong, significant correlation between fungal and bacterial community structure of individual samples at the level of ASVs using Procrustes rotation (cor. = 0.29, p < 0.001; Fig. 2C). Collapsing ASV taxonomy to genus, family, and order resulted in even stronger correlations (cor. = 0.44, 0.48 & 0.43, respectively; all p < 0.001; Fig. 2C). These data indicate a coupling between the structures of fungal and bacterial communities, whereby shifts in structure of one community across the phylogeny also reflect consistent shifts in the other microbial group.

### Strength of Interactions Between Bacteria and Fungi May Vary Across Host Taxonomy

Analysis of correlations among fungal and bacterial abundances revealed differences in network structure at both the host class (Fig. 3A) and host species level (Figs. S7; S8). In particular, fungi of the phylum Ascomycota appeared frequently in the putative interaction networks of birds, mammals and amphibians (Fig. 3A). There was also systematic variation in network structure among taxonomic groups. Using the class-level network data in Fig. 3A, we estimated that Mammalia exhibited the fewest components, fewest communities, and lowest modularity (Table 2), indicating lower overall network subdivision relative to other animal classes. Mean betweenness of fungal nodes also varied by host class; randomisations revealed that mean fungal betweenness was significantly lower than expected by chance in Aves (2-tailed p = 0.044, Fig. 3B) but not Mammalia (2-tailed p=0.6, Fig 3B). Models of species-level network data (Fig. S7, S8) revealed the frequency of positive co-occurrence between pairs of microbes also varied by class; Mammalia exhibited the highest proportion of positive edges (Fig. 3C), being significantly greater than those of birds (mean diff. 0.042 [0.017-0.067]) and amphibians (mean diff. 0.05 [0.002-0.112]). Notably, insects had a markedly lower proportion of positive edges compared to all other taxa (Fig. 3C). Class explained 93.2% [92.9-93.4%] of variation in edge sign.

**FIGURE 3.**
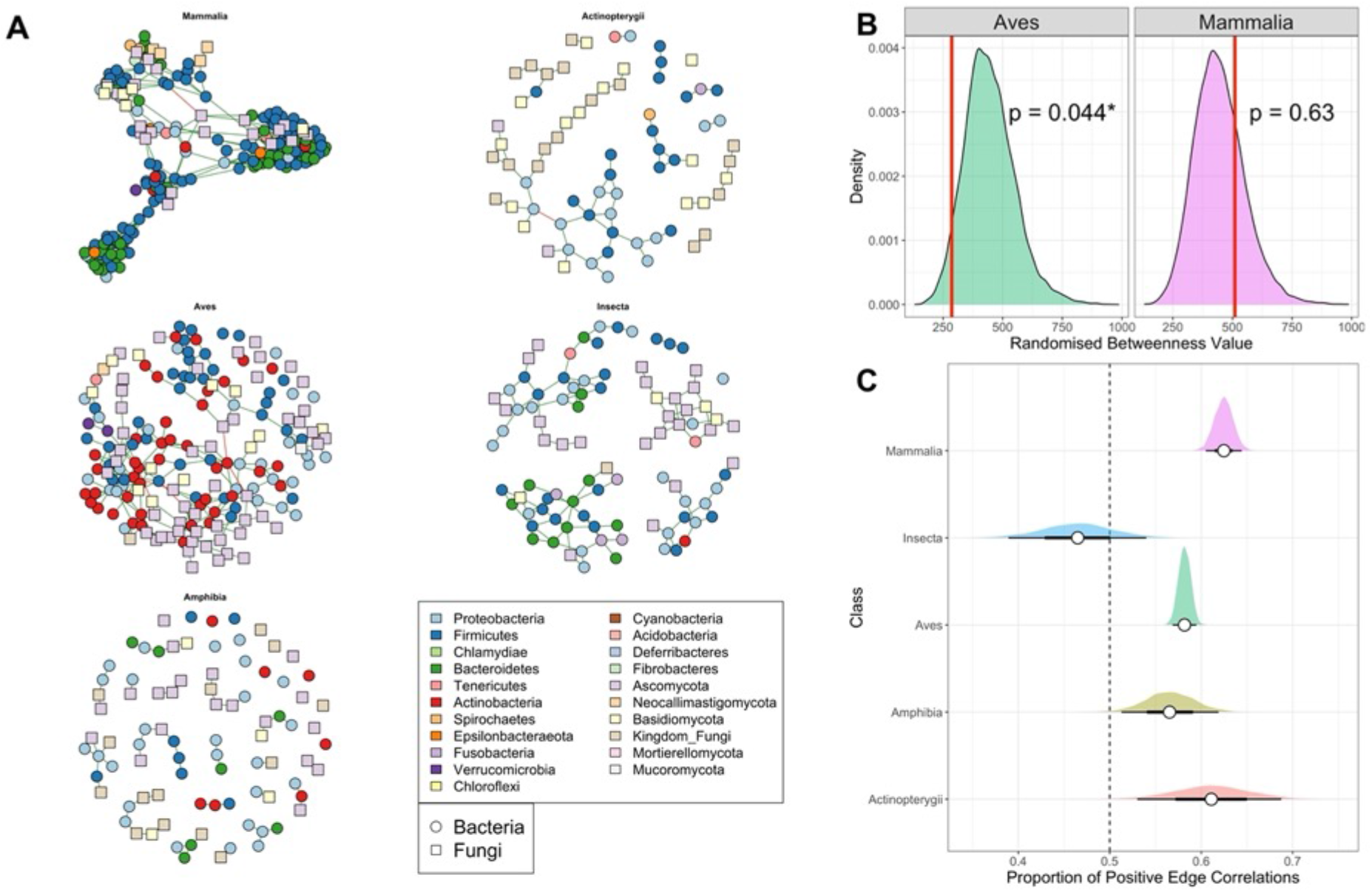
**(A)** Putative microbial interaction networks between bacterial (circles) and fungal (squares) taxa, coloured by microbial phylum. Networks were constructed using the R package *SpiecEasi* on CLR-transformed abundance values to detect non-random co-occurrence between groups of microbes. **(B)** Permutational testing revealed that mean fungal betweenness was significantly lower than expected by chance in Aves, but not Mammalia, indicating heterogeneity in network structure. **(C)** Analysis of network structural traits from species-specific networks comprising 39 species from five Classes. There were significant differences in the proportion of positive edges (correlations between paired microbial abundance values) among classes. Vertical dashed line indicates equal proportion of positive and negative edges.

## DISCUSSION

Our study represents the most wide-ranging evaluation of animal mycobiome composition, and its covariation with the bacterial microbiome, undertaken to date. Our data provide novel evidence for mycobiome phylosymbiosis in wild animals, indicative of close evolutionary coupling between hosts and their resident fungal communities. Consistent with previous studies, we also find evidence of phylosymbiosis in the bacterial microbiome (29), but crucially, we demonstrate strong and consistent covariation between fungal and bacterial communities across host phylogeny, especially at higher microbial taxonomic levels. These patterns are supported by complementary network analysis illustrating frequent correlative links between fungal and bacterial taxa, whereby certain pairs of microbes from different kingdoms are much more likely to co-occur in the microbiome than expected by chance. Taken together, these data provide novel evidence of host recruitment for specific fungal and bacterial communities, which in turn may reflect host selection for interactions between bacteria and fungi critical for host physiology and health.

We found marked variation among host species in microbial community richness and composition for both bacteria and fungi. Though our data suggest many species support a diverse assemblage of host-associated fungi, we show that bacterial diversity tends to be higher on average relative to fungal diversity, and that there is no signal of positive covariance between fungal and bacterial richness within species, suggesting more ASV-rich bacterial microbiomes are not consistently associated with more ASV-rich mycobiomes. These patterns could arise because of competition for niche space within the gut, where high bacterial diversity may reflect stronger competition that prevents proliferation of fungal diversity. Understanding patterns of niche competition within and among microbial groups requires that we are able to define those niches by measuring microbial gene function, and quantifying degree of overlap or redundancy in functional genomic profiles across bacteria and fungi.

We detected strong phylosymbiosis for both fungi and bacteria across a broad host phylogeny encompassing both vertebrate and invertebrate classes. This pattern was significantly stronger in bacteria than for fungi. In both microbial kingdoms, the signal of phylosymbiosis strengthened when aggregating microbial assignments to broader taxonomic levels, a phenomenon that has previously been shown for bacterial communities (30). That this pattern also occurs in fungi suggests either that host recruitment is weaker at finer-scale taxonomies, or our ability to detect that signal is weaker at the relatively noisy taxonomic scale of ASVs. Stronger signals of phylosymbiosis at family and order-level taxonomies may reflect the deep evolutionary relationships between hosts and their bacterial and fungal communities, as well as the propensity for microbial communities to allow closely related microbes to establish whilst repelling less related organisms (31). That is, higher-order microbial taxonomy may better approximate functional guilds within the microbiome, such as the ability to degrade cellulose (25, 30), which are otherwise obscured by taxonomic patterns of ASVs. Resolving this requires the integration of functional genomic data from the fungal and bacterial microbiota into the phylogeny.

In addition to microbe-specific patterns of phylosymbiosis, a key novel finding of our work is discovery of strong covariation between fungal and bacterial community composition across the host phylogeny. These patterns are consistent with host recruitment for particular suites of fungal and bacterial taxa, which may represent bacteria-fungi metabolic interactions beneficial to the host. Bacterial-fungal interactions have previously been demonstrated for a handful of animal species (8, 9, 17, 32, 33), but here we show these are widespread across multiple animal classes. Both bacteria and fungi have considerable enzymatic properties that facilitate the liberation of nutrients for use by other microbes, thus facilitating cross-kingdom colonisation (34–36) and promoting metabolic inter-dependencies (37–39). We also identified numerous associations between bacteria and fungi for many of our host species. The frequency and predicted direction of these relationships varied considerably among host classes, with the mammalian network exhibiting i) a lower modularity, indicating weaker clustering into fewer discrete units (both distinct components and interlinked communities); and ii) a higher frequency of positive correlations between microbes compared to most other classes, in particular birds and insects. Comparisons of networks are challenging when they differ in size (i.e., number of nodes) and structure, and differences between classes in traits like modularity will also be affected by species replication within each class. However, proportional traits like interaction structure (proportion of positive interactions) are unlikely to be driven solely by sample size, suggesting marked biological variation in strength of fungi-bacteria interactions across the host phylogeny. These putative interaction networks provide novel candidates for further investigation in controlled systems, where microbiome composition and therefore the interactions among microbes can be manipulated to test the influence of such interactions on host physiology.

The drivers of phylosymbiosis remain unclear, even for bacterial communities; is a phylogenetic signal indicative of host-microbiome coevolution, or simply a product of “ecological filtering” of the microbiome in the host organism either via extrinsic (e.g. diet, habitat) or intrinsic sources (e.g. gut pH, immune system function) (26, 29, 40)? Our results indicate host diet may play a role in determining bacterial composition in mammals, but not fungal composition in either mammals or birds. These results are broadly consistent with previous work, where the influence of diet on bacterial microbiome was most evident in mammals (25). However, Li et al. (16) showed that the composition and diversity of both fungal and bacterial communities of faecal samples differed between phytophagous and insectivorous bats, and Heisel et al. (17) demonstrated changes in fungal community composition in mice fed a high fat diet. Our study was not designed to test for the effects of ecological variation in diet on fungal microbiome *within* a species, nor can we discount the possibility that at finer taxonomic scales within classes, signals of the effect of *among* species variation in diet on mycobiome may become stronger (e.g. (16)). It is also worth noting that the signals produced from faecal and true gut samples may differ; evidence suggests faecal samples may indicate diet is the predominant driver of “gut” microbiome composition when gastrointestinal samples indicate host species is the predominant determinant (41). Moreover, faecal samples may only represent a small proportion of the gastrointestinal microbiome (41–43). Our data also show that sample type has a significant effect on both fungal and bacterial community composition (as well as DNA extraction method and storage method; see (44–47) for other examples of this). As such, a more thorough analysis of true gut communities is required to determine the extent to which mycobiome phylosymbiosis and dietary signals occur across wild animals, and what other ecological and host-associated factors influence mycobiome composition and function. We hypothesise that evolutionary processes play a large role in shaping host-associated microbiomes, with selection for microbiome function rather than taxonomic groupings per se.

Within animals, the roles of host-associated fungal communities are not well understood, yet our data highlight that fungi are important components of microbiome structure that are often overlooked. Our knowledge of the range of functions provided by the host mycobiome, and how these alter or complement those provided by the bacterial microbiome, remains limited. We hypothesise that host-associated fungi and bacteria produce mutually beneficial metabolites that facilitate the colonisation, reproduction and function of cross-kingdom metabolic networks (28). Though we provide evidence for consistent variation among host class in fungal community structure, and the role of fungi within putative interaction networks, for many researchers the questions of key interest will focus on what governs variation at the level of the individual. Clear gaps in our knowledge remain regarding the relative contributions of host genomic (48–50) and environmental variation to host mycobiome structure, function and stability. We argue that there is an urgent need to incorporate quantitative estimates of microbial function into microbiome studies, which are crucial for understanding the forces of selection shaping host-microbe interactions at both the individual and species level.

## MATERIALS AND METHODS

### Sample collection

DNA was extracted from tissue or faecal samples of 49 host species using a variety of DNA extraction methods (Table S1) and normalised to ~10 ng/ul. Samples were largely collated from previous studies and/or those available from numerous researchers and as such, DNA extraction and storage techniques were not standardised across species. We sequenced a median of 10 samples per species (range of 5 to 12; Table S1).

### ITS1F-2 and 16S rRNA amplicon sequencing

Full details are provided in Supplementary Materials. Briefly, we amplified the ITS1F-2 rRNA gene to identify fungal communities using single index reverse primers and a modified protocol of Smith & Peay (51) and Nguyen et al. (52), as detailed in Griffiths et al. (13). To identify bacterial communities, we amplified DNA for the 16S rRNA V4 region using dual indexed forward and reverse primers according to Kozich et al. (53) and Griffiths et al. (49). The two libraries were sequenced separately using paired-end reads (2 x 250bp) with v2 chemistry on an Illumina MiSeq.

We conducted amplicon sequence data processing in DADA2 v1.5 (54) in RStudio v1.2.1335 for R (55, 56) for both ITS rRNA and 16S rRNA amplicon data. After data processing, we obtained a median of 1425 reads per sample (range of 153 to 424,527) from the ITS data, and a median of 3273 reads (range of 153 to 425,179) for the 16S rRNA data.

To compare alpha-diversity between species and microbial kingdoms, we rarefied libraries to 500 reads per sample, yielding 292 samples from 46 species and 307 samples from 47 species for fungal and bacterial kingdoms respectively. Alpha-diversity measures remained relatively stable within a host species whether data were rarefied to 500, 1000, or 2500 reads (Figs. 1, S1, S2; see Supplementary Material for more details).

### Host phylogeny

As many of our host species lack genomic resources from which to construct a genome-based phylogeny, we built a dated phylogeny of host species using TimeTree (57). The phylogenetic tree contained 42 species, of which 36 were directly represented in the TimeTree database. A further six species had no direct match in TimeTree and so we used a congener as a substitute (*Amietia, Glossina, Portunus, Ircinia, Amblyomma, Cinachyrella*). We calculated patristic distance among species based on shared branch length in the phylogeny using the ‘cophenetic’ function in the *ape* package (58) in R. We visualised and annotated the phylogeny using the R package *ggtree* (59). To create a phylogeny for all samples, we grafted sample-level tips onto the species phylogeny with negligible branch lengths following Youngblut et al. (25).

### Fungal and bacterial community analysis

A fully reproducible workflow of all analyses is provided in supplementary material as an R Markdown document. We used the R package *brms* (60, 61) to fit (generalized) linear mixed effects models [(G)LMMs] to test for differences in alpha diversity and calculated r^2^ of models using the ‘bayes_R2’ function. We assessed the importance of terms based on whether 95% credible intervals of the parameter estimates of interest crossed zero. We used *ggplot* (62), *cowplot* (63) and *tidybayes* (64) for raw data and plotting of posterior model estimates.

To support these analyses, we also used the R packages *phylobase* (65) and *phylosignal* (66) to estimate the phylogenetic signal in patterns of alpha diversity for both bacteria and fungi, using both Inverse Simpson Index and number of observed ASVs as outcome variables. We calculated Abouheif’s C_mean_ for each diversity-microbe combination and corrected p values for multiple testing using Benjamini-Hochberg correction.

To identify taxonomic differences in microbiome and mycobiome composition between host species, we used centred-log-ratio (CLR) transformation in the *microbiome* (67) package to normalise microbial abundance data, which obviates the need to lose data through rarefying (68). To quantify differences in beta-diversity among kingdoms and species whilst simultaneously accounting for sample storage and library preparation differences among samples, we conducted a PERMANOVA analysis on among-sample Euclidean distances of CLR-transformed abundances using the *adonis* function in *vegan* (69) with 999 permutations. For both kingdoms, we specified effects in the following order: sample type, tissue storage, extraction kit, class, order, species. This marginalises the effects of sample metadata variables first, before partitioning the remaining variance into that accounted for by host phylogeny. The results were similar when amplicon data were converted to relative abundance or rarefied to 500 reads (data not presented).

To test the hypothesis that inter-individual differences in microbial community composition were preserved between microbial kingdoms, we performed Procrustes rotation of the two PCA ordinations for bacterial and fungal abundance matrices, respectively (n = 277 paired samples from 46 species). We also repeated this analysis with ASVs agglomerated into progressively higher taxonomic rankings from genus to order (see (30)). To provide a formal test of differences in strength of covariation at different taxonomic levels, we conducted a bootstrap resampling analysis where for each kingdom at each iteration, we randomly sampled 90% of the data and recalculated the correlation metric. We repeated this process 999 times to build a distribution of correlation values at each taxonomic grouping. To examine the hypothesis that inter-individual distance in microbial community composition varies in concert with interspecific phylogenetic distance, we performed a Procrustes rotation on the paired matrix of microbial distance (Euclidean distance of CLR-transformed abundances) and patristic distance from the phylogenetic tree.

To identify potential co-occurrence relationships between fungal and bacterial communities, we conducted two analyses; 1) We used the R package *SpiecEasi* (70) to identify correlations between unrarefied, CLR-transformed ASVs abundances at the host class level (with insects grouped), and 2) we used co-occurrence analysis at the species level, by rarefying the bacterial and fungal data sets to 500 reads each, and agglomerated taxonomy family level, resulting in 117 bacterial groups and 110 fungal groups. We then merged the *phyloseq* objects for bacterial and fungal communities for each sample, with sufficient data retained to conduct the co-occurrence analysis for 40 host species. Using these cross-kingdom data, we calculated the co-occurrence between each pair of microbial genera by constructing a Spearman’s correlation coefficient matrix in the *bioDist* package (71, 72). We visualised those with rho > 0.50 (strong positive interactions) and rho < −0.50 (strong negative interactions) for each host species separately using network plots produced in *igraph* (73). We calculated modularity of the class-level microbial networks comprising both positive and negative interactions using the modularity function after greedy clustering implemented in the *igraph* package. We used binomial GLM to test the hypothesis that the proportion of positive edges (correlations) varies by host class, and permutation approaches on betweenness values of fungal nodes to test the hypothesis that fungi form critical components of microbial networks.

To determine the effect of diet on bacterial and fungal community composition, we used only samples from the bird and mammal species and agglomerated the data for each host species using the merge_samples function in *phyloseq* (74). This gave us a representative microbiome for each host species, which we rarefied to the lowest number of reads for each combination of kingdom and host taxon (2,916 – 9,160 reads; bacterial read counts were low for lesser horseshoe bats and so this species was removed from this analysis) and extracted Euclidean distance matrices for each. We then correlated these with dietary data obtained from the EltonTraits database (75) using Mantel tests with Kendall rank correlations in the *vegan* package (69). We agglomerated the microbial data to class level and visualised the bacterial and fungal community compositions for mammals alongside pie charts displaying EltonTrait dietary data for each species. We also used a primary axis of the ordination of EltonTrait data to derive a ‘dietary variation axis’ used as a predictor for alpha diversity of Birds and Mammals.

## ACKNOWLEDGEMENTS

We would like to thank Miran Aprahamian, Chris Williams (Environment Agency) Patrick Abila (National Livestock Resources Research Institute) and Dr Patrick Vudriko (Makerere University) for providing samples, as well as BEI Resources, the US Forest Service and the University of Wisconsin-Madison for providing mock communities. We are grateful to Devenish Nutrition for funding the red deer research at Dowth Hall, Co. Meath, Ireland. The sampling of shrews was funded by a Heredity Fieldwork Grant awarded by the Genetics Society. The collection of dove and pigeon faecal samples was jointly funded by the Royal Society for the Protection of Birds and Natural England through the Action for Birds in England (AfBiE) partnership. Fieldwork enabling collection of avian faecal samples from Lincolnshire was funded by The Royal Society Research Grant RG170086 to JCD. Small mammal sampling in the Chernobyl Exclusion Zone was supported by the TREE (https://tree.ceh.ac.uk/) and RED FIRE (https://www.ceh.ac.uk/redfire) projects. TREE was funded by the Natural Environment Research Council (NERC), Radioactive Waste Management Ltd. and the Environment Agency as part of the RATE Programme; RED FIRE was a NERC Urgency Grant. Northern muriqui research at Caparaó National Park, Brazil, was funded by CAPES (BEX 1298/15-1).

## COMPETING INTERESTS

The authors have no competing interests to declare.

## DATA ACCESSIBILITY STATEMENT

Sequence data are deposited in the NCBI SRA database under BioProject numbers PRJNA593927 and PRJNA593220. A fully-reproducible analysis workflow has been provided as supplementary material at https://github.com/xavharrison/Mycobiome2020

## Notes

### Competing Interest Statement

The authors have declared no competing interest.

### Summary of Updates

Figure 1 and main text updated to include host dietary analysis; Figure 2 updated to include additional analysis on host-microbe and bacteria-fungi correlations; Figure 3 and 4 merged and updated to include randomisations of network structure; main text streamlined.

