## Supplementary Tables and Figures for "Fungal microbiomes are determined by host phylogeny and exhibit widespread associations with the bacterial microbiome"

### METHODS

*ITS1F-2 and 16S rRNA amplicon sequencing*

To identify fungal communities, we amplified DNA for the ITS1F-2 rRNA gene using single index reverse primers and a modified protocol of Smith & Peay (1) and Nguyen et al. (2), as detailed in Griffiths et al. (3). We ran PCRs in duplicate using Solis BioDyne 5x HOT FIREPol® Blend Master Mix, 2µM primers and 1.5µl of sample DNA. Thermocycling conditions were 95 °C for 10 min, followed by 30 cycles of 95 °C for 30s, 52 °C for 20s and 72 °C for 30s, with a final extension of 72 °C for 8 minutes. We quality checked the PCR products using a 2200 TapeStation (Agilent, USA). We combined PCR replicates into a single PCR plate and cleaned products using HighPrep™ PCR clean up beads (MagBio, USA) according to the manufacturers' instructions. To normalise the libraries, we combined 1ul of each sample and conducted a titration sequencing run with this pool using an Illumina v2 nano cartridge (paired end reads; 2 x 150bp) on the Illumina MiSeq at the University of Salford. Based on the percentage of reads sequenced per library, we calculated the volume required for the full sequencing run and pooled these accordingly. ITS rRNA amplicon sequencing was conducted using paired-end reads (2 x 250bp) using an Illumina v2 cartridge on the MiSeq platform at the University of Salford. We included negative controls (blank extractions) for six of the nine DNA extraction methods plus a blank consisting of PCR-grade water, as well as a fungal mock community as a positive control. We ran the same library twice to increase sequencing depth, and combined data within samples across these two runs in the data pre-processing stage.

To identify bacterial communities, we amplified DNA for the 16S rRNA V4 region using dual indexed forward and reverse primers according to Kozich et al. (4) and Griffiths et al. (5). We ran PCRs in duplicate as described above using thermocycling conditions of 95°C for 15 minutes, followed by 28 cycles of 95°C for 20s, 50°C for 60s and 72°C for 60s, and a final extension at 72°C for 10 minutes. After cleaning and quality checking (as above), we again sequenced an equivolume pool on an Illumina v2 nano cartridge as described above, then pooled samples according to read coverage and conducted a full paired-end sequencing run (2 x 250bp) using Illumina v2 chemistry. We included extraction blanks and a mock bacterial community as negative and positive controls, respectively.

*Pre-processing of amplicon sequence data*

We conducted all data processing and analysis in RStudio v1.2.1335 for R (6, 7) (see supplementary files for full code). For 16S rRNA amplicon sequencing data, adapters and primers were automatically trimmed by the MiSeq BaseSpace software, but for ITS rRNA amplicon data we performed an additional trimming step in cutadapt (8) to remove these. We conducted amplicon sequence processing in DADA2 v1.5 (9) for both ITS rRNA and 16S rRNA amplicon data.

A total of 8,033,962 raw sequence reads from 934 samples (i.e. duplicate data for each sample from the two sequencing runs) were generated across the two ITS rRNA sequencing runs. Modal contig length was 225 bp (range 112-477 bp) once paired-end reads were merged. We did not conduct additional trimming based on sequence length as the ITS region is highly variable (10). We removed 29 amplicon sequence variants (ASVs) found in the negative controls and filtered out chimeras, and then assigned taxonomy using the UNITE v7.2 database (11). We combined sequence data for each sample across the two ITS rRNA sequencing runs using the merge_samples function in phyloseq (12). After data processing, we obtained a median of 1425 reads per sample (range of 153 to 424,527). DADA2 identified 12 unique ASVs in the sequenced mock community sample comprising 12 fungal isolates.

A total of 6,657,351 raw sequence reads from 476 samples were generated during 16S rRNA sequencing. Modal contig length was 253 bp once paired-end reads were merged. We removed ASVs with length >260 bp (55 SVs; 0.004% of total sequences) along with chimeras and 17 SVs found in the negative controls. We assigned taxonomy using the SILVA v132 database (13, 14). We stripped out chloroplasts and mitochondria from samples, leaving a median of 3273 reads per sample (range of 153 to 425,179). DADA2 identified 20 unique ASVs in the sequenced mock community sample comprising 20 bacterial isolates.

*Alpha Diversity Models*

For model fitting, we filtered the data to only those samples with paired metrics of microbial richness for both kingdoms (201 observations from 42 species). For higher order taxonomic predictors and random effects, we binned all invertebrate classes into a single grouping to improve model performance, as otherwise invertebrate class and species were co-linear. All vertebrate taxonomic groupings were equivalent to class (Mammalia, Aves etc). We fitted two models to these data. First, to quantify relative differences in richness between bacteria and fungi within a sample, we used GLMMs in the *brms* package, with i) Bernoulli errors and a logit link; ii) a binary response of '1' if bacterial richness was higher than fungal richness, and '0' otherwise; and iii) 'Species' nested within 'Class' as random intercepts. We did not include intermediate levels of taxonomy because replication at Order and Family levels was low relative to Class. We did not use a phylogenetic mixed model as not all species were represented in the TimeTree phylogeny. Second, to quantify absolute differences in microbial richness, we fitted a bivariate response LMM with both fungal and bacterial richness values as a two-column response with Class as a fixed effect, and Species as a random intercept. For all models, we used uninformative Cauchy priors for the random effects and Gaussian priors for fixed effects coefficients. We assessed model adequacy using visual inspection of chains to assess mixing and stationarity properties, as well as posterior predictive checks using the 'pp_check' function in *brms*.

*Beta Diversity Analysis*

To visualise differences in microbial community structure among samples, we i) plotted proportional abundance of microbial groups at the phylum level, aligned to the host phylogenetic tree, ii) agglomerated the data to class level and visualised the variation in CLR-transformed ratios for the five most abundant microbial classes in each kingdom for each species using jitter plots, and iii) conducted principal components analysis (PCA) using CLR-transformed abundance matrices for each kingdom

**TABLE S1:** Details of host species and their origins, sex ratios, sample sizes and types, and storage and extraction methods for the study.

| Class | Common name | Scientific name | N | Sex ratio (M: F: J: unknown: N/A) | Captive or Wild | Origin | Sample Type | Collection Year | Tissue Storage | Extraction Kit |
|---|---|---|---|---|---|---|---|---|---|---|
| Demospongia | Vase sponge | *Ircinia campana* | 10 | 0: 0: 0: 0: 10 | Wild | Long Key, Florida, USA | Tissue (choanosome) | 2014 | 95% ethanol | Qiagen Blood and Tissue kit with proteinase K |
| Demospongia | Golfball sponge | *Cinachyrella sp.* | 10 | 0: 0: 0: 0: 10 | Wild | Long Key, Florida, USA | Tissue (choanosome) | 2014 | 95% ethanol | Qiagen Blood and Tissue kit with proteinase K |
| Arachnida | Hard tick | *Amblyomma rotundatum* | 10 | 0: 0: 0: 10: 0 | Wild | Montserrat, Caribbean | Whole organism | 2014 | 70% ethanol | Alkaline digest and ethanol precipitation |
| Malacostraca | Blue swimming crab | *Portunus segnis* | 5 | 0: 0: 0: 5: 0 | Wild | Malta | Gut | 2018 | 70% ethanol | Quigen QIAamp Fast DNA Stool Mini kit |
| Malacostraca | Brown shrimp | *Crangon crangon* | 10 | 0: 0: 0: 10: 0 | Wild | Liverpool, Lancashire, England | Gut | 2018 | Buffer AE and frozen at -20°C | Qiagen Blood and Tissue kit with proteinase K |
| Insecta | Cockroach | *Diploptera punctata* | 11 | 7: 1: 0: 3: 0 | Captive | Manchester Metropolitan University, Manchester, UK | Gut | 2018 | Liquid nitrogen and frozen at -80°C | Qiagen Blood and Tissue kit with proteinase K and lysozyme |
| Insecta | Honey bee | *Apis mellifera* | 10 | 0: 10: 0: 0: 0 | Wild | North West of England, UK | Gut | 2016 | 100% ethanol and frozen at -20°C | Qiagen Blood and Tissue kit with proteinase K and lysozyme |
| Insecta | Tsetse fly | *Glossina fuscipes* | 9 | 2: 7: 0: 0: 0 | Wild | Patira East, Uganda | Whole organism | 2019 | 70% ethanol | Qiagen Blood and Tissue kit with proteinase K |
| Insecta | African palm weevil larvae | *Rhynchophorus phoenicis* | 6 | 0: 0: 0: 6: 0 | Wild | Sapele Town, Delta State, Nigeria | Gut | 2019 | Frozen at -20°C | ZymoBIOMICS DNA mini kit |
| Actinopterygii | European eel | *Anguilla anguilla* | 10 | 0: 0: 0: 10: 0 | Wild | Cumbria, England | Gut | 2009 | Frozen at -20°C | Qiagen PowerSoil kit |
| Actinopterygii | Foureye butterflyfish | *Chaetodon capistratus* | 10 | 0: 0: 0: 10: 0 | Wild | Bocas del Toro, Bahia Almirante, Panama | Gut | 2018 | 95% ethanol | Qiagen PowerSoil kit with proteinase K |
| Actinopterygii | Yellowhead wrasse | *Halichoeres garnoti* | 10 | 0: 0: 0: 10: 0 | Wild | Caye Caulker, Belize | Gut | 2015 | 95% ethanol | Qiagen PowerSoil kit with proteinase K |
| Actinopterygii | Barred hamlet | *Hypoplectrus puella* | 12 | 0: 0: 0: 12: 0 | Wild | Bocas del Toro, Bahia Almirante, Panama | Gut | 2018 | 95% ethanol | Qiagen PowerSoil kit with proteinase K |
| Amphibia | Common midwife toad | *Alytes obstetricans* | 11 | 0: 0: 0: 11: 0 | Captive | London Zoo, London, UK | Skin swab | 2015 | Frozen at -20°C | Qiagen DNEasy kit |

| | | | | | | | | | | |
|---|---|---|---|---|---|---|---|---|---|---|
| Amphibia | Phofung river frog | *Amietia hymenopus* | 10 | 0: 0: 10: 0: 0 | Wild | Drakensberg National Park, South Africa | Tadpole mouthparts | 2015 | 95% ethanol | Qiagen Blood and Tissue kit with proteinase K |
| Amphibia | Common toad | *Bufo bufo* | 10 | 0: 0: 10: 0: 0 | Wild | Norway | Whole organism | 2009 | 70% ethanol | Phenol chlorophorm |
| Amphibia | Great-crested newt | *Triturus cristatus* | 10 | 5: 5: 0: 0: 0 | Wild | Lancashire, England | Toe clip | 2015 | 70% ethanol | Phenol chlorophorm |
| Aves | Reed warbler | *Acrocephalus scirpaceus* | 8 | 3: 3: 0: 2: 0 | Wild | Lincolnshire, UK | Faeces | 2018/19 | Frozen at -20°C | Qiagen PowerSoil Pro kit |
| Aves | Light-bellied brent goose | *Branta bernicla hrota* | 10 | 0: 0: 0: 10: 0 | Wild | Iceland | Faeces | 2017 | Frozen at -20°C | Qiagen PowerSoil kit |
| Aves | Goldfinch | *Carduelis carduelis* | 8 | 5: 1: 0: 2: 0 | Wild | Lincolnshire, UK | Faeces | 2018/19 | Frozen at -20°C | Qiagen PowerSoil Pro kit |
| Aves | Stock dove | *Columba oenas* | 10 | 0: 0: 0: 10: 0 | Wild | East Anglia, UK | Faeces | 2014 | Frozen at -20°C | Qiagen QIAamp Fast DNA Stool Mini kit |
| Aves | Woodpigeon | *Columba palumbus* | 5 | 0: 0: 0: 5: 0 | Wild | East Anglia, UK | Faeces | 2012 | Frozen at -20°C | Qiagen QIAamp Fast DNA Stool Mini kit |
| Aves | Carrion crow | *Corvus corone* | 12 | 0: 0: 0: 12: 0 | Wild | Cumbria, UK | Gut | 2019 | Frozen at -20°C | Qiagen Microbiome kit |
| Aves | Blue tit | *Cyanistes caeruleus* | 8 | 0: 0: 0: 8: 0 | Wild | Lincolnshire, UK | Faeces | 2018 | Frozen at -20°C | Qiagen PowerSoil Pro kit |
| Aves | Yellowhammer | *Emberiza citrinella* | 8 | 1: 1: 0: 6: 0 | Wild | Lincolnshire, UK | Faeces | 2018 | Frozen at -20°C | Qiagen PowerSoil Pro kit |
| Aves | Reed bunting | *Emberiza schoeniclus* | 8 | 4: 3: 0: 1: 0 | Wild | Lincolnshire, UK | Faeces | 2018 | Frozen at -20°C | Qiagen PowerSoil Pro kit |
| Aves | Robin | *Erithacus rubecula* | 8 | 1: 1: 0: 6: 0 | Wild | Lincolnshire, UK | Faeces | 2018 | Frozen at -20°C | Qiagen PowerSoil Pro kit |
| Aves | Great tit | *Parus major* | 8 | 3: 3: 0: 2: 0 | Wild | Lincolnshire, UK | Faeces | 2018/19 | Frozen at -20°C | Qiagen PowerSoil Pro kit |
| Aves | Chiffchaff | *Phylloscopus collybita* | 8 | 0: 1: 0: 7: 0 | Wild | Lincolnshire, UK | Faeces | 2018/19 | Frozen at -20°C | Qiagen PowerSoil Pro kit |
| Aves | Collared dove | *Streptopelia decaocto* | 8 | 0: 0: 0: 8: 0 | Wild | East Anglia, UK | Faeces | 2014 | Frozen at -20°C | Qiagen QIAamp Fast DNA Stool Mini kit |
| Aves | Turtle dove | *Streptopelia turtur* | 7 | 0: 0: 0: 7: 0 | Wild | East Anglia, UK | Faeces | 2014 | Frozen at -20°C | Quigen QIAamp Fast DNA Stool Mini kit |
| Aves | Blackcap | *Sylvia atricapilla* | 8 | 3: 2: 0: 3: 0 | Wild | Lincolnshire, UK | Faeces | 2018 | Frozen at -20°C | Qiagen PowerSoil Pro kit |
| Aves | Song thrush | *Turdus philomelos* | 8 | 0: 0: 0: 8: 0 | Wild | Lincolnshire, UK | Faeces | 2018 | Frozen at -20°C | Qiagen PowerSoil Pro kit |
| Mammalia | Striped field mouse | *Apodemus agrarius* | 10 | 5: 5: 0: 0: 0 | Wild | Chernobyl Exclusion Zone, Ukraine | Faeces | 2017 | 100% ethanol and frozen at -20°C | Invitrogen Microbiome kit |

| | | | | | | | | | | |
|---|---|---|---|---|---|---|---|---|---|---|
| Mammalia | Yellow-necked mouse | *Apodemus flavicollis* | 10 | 5: 5: 0: 0: 0 | Wild | Chernobyl Exclusion Zone, Ukraine | Faeces | 2017 | 100% ethanol and frozen at -20°C | Invitrogen Microbiome kit |
| Mammalia | Wood mouse | *Apodemus sylvaticus* | 10 | 6: 4: 0: 0: 0 | Wild | Chernobyl Exclusion Zone, Ukraine | Faeces | 2017 | 100% ethanol and frozen at -20°C | Invitrogen Microbiome kit |
| Mammalia | Northern muriqui | *Brachyteles hypoxanthus* | 10 | 0: 0: 0: 10: 0 | Wild | Caparao National Park, Espirito Santo, Brazil | Faeces | 2017/18 | RNA Later and frozen at -20°C | Qiagen QIAamp Fast DNA Stool Mini kit |
| Mammalia | Roe deer | *Capreolus capreolus* | 7 | 7: 0: 0: 0: 0 | Wild | Cumbria, UK | Faeces | 2019 | Frozen at -20°C | Qiagen Microbiome kit |
| Mammalia | Red deer | *Cervus elaphus* | 10 | 0: 0: 0: 10: 0 | Wild | County Meath, Ireland | Faeces | 2018 | Frozen at -20°C | Qiagen QIAamp Fast DNA Stool Mini kit |
| Mammalia | Greater white-toothed shrew | *Crocidura russula* | 10 | 5: 5: 0: 0: 0 | Wild | Belle Ile, France | Gut | 2018 | 100% ethanol and frozen at -20°C | Qiagen PowerSoil kit |
| Mammalia | Eastern black rhino | *Diceros bicornis michaeli* | 10 | 0: 10: 0: 0: 0 | Captive | Chester Zoo and Port Lympne Wild Animal Park, UK | Faeces | 2011 | Frozen at -20°C | Qiagen QIAamp Fast DNA Stool Mini kit |
| Mammalia | Wild pony | *Equus ferus caballus* | 10 | 5: 5: 0: 0: 0 | Wild | Snowdonia National Park, Wales | Faeces | 2013 | Frozen at -20°C | Qiagen QIAamp Fast DNA Stool Mini kit |
| Mammalia | Hedgehog | *Erinaceus europaeus* | 12 | 0: 0: 0: 12: 0 | Wild | Cumbria, UK | Faeces | 2019 | Frozen at -20°C | Qiagen Microbiome kit |
| Mammalia | Bank vole | *Myodes glareolus* | 10 | 7: 3: 0: 0: 0 | Wild | Chernobyl, Ukraine | Gut | 2017 | 100% ethanol and frozen at -20°C | Invitrogen Microbiome kit |
| Mammalia | Lesser horseshoe bat | *Rhinolophus hipposideros* | 10 | 3: 5: 2: 0: 0 | Wild | County Kerry, Ireland | Faeces | 2016 | Frozen at -20°C | Zymo DNA Extraction kit |
| Mammalia | Capuchin monkey | *Sapajus libidinosus* | 10 | 0: 0: 0: 10: 0 | Wild | Serra Talhada, State of Pernambuco/Minas Gerais, Brazil | Faeces | 2017 | Frozen at -20°C | Qiagen QIAamp Fast DNA Stool Mini kit |
| Mammalia | Grey squirrel | *Sciurus carolinensis* | 12 | 0: 0: 0: 12: 0 | Wild | Cumbria, UK | Faeces | 2019 | Frozen at -20°C | Qiagen Microbiome kit |
| Mammalia | Red squirrel | *Sciurus vulgaris* | 12 | 0: 0: 0: 12: 0 | Wild | Cumbria, UK | Faeces | 2019 | Frozen at -20°C | Qiagen Microbiome kit |
| Mammalia | Pygmy shrew | *Sorex minutus* | 10 | 5: 5: 0: 0: 0 | Wild | Belle Ile, France | Gut | 2018 | 100% ethanol and frozen at -20°C | Qiagen PowerSoil kit |

### RESULTS

Alpha-diversity measures remained relatively stable within a host species whether data were rarefied to 500, 1000, or 2500 reads (Figures 1, S1, S2; Supplementary Material). Patterns between kingdoms were similar for each host species whether data were rarefied to 500 or 1000 reads, with the exception of slight increases in fungal diversity relative to bacterial diversity for two host species (blue tit, light-bellied brent goose) when data were rarefied to 1000 reads (Figure S1). Cross-kingdom patterns for each host species were also similar whether data were rarefied to 500 or 2500 reads (Figure S2), although four host species (chiffchaff, greater white-toothed shrew, light-bellied brent goose, reed warbler) showed greater differences between bacterial and fungal diversity when 2500 reads were used, and one (common midwife toad) had reduced differences (Figures 1 and S2).

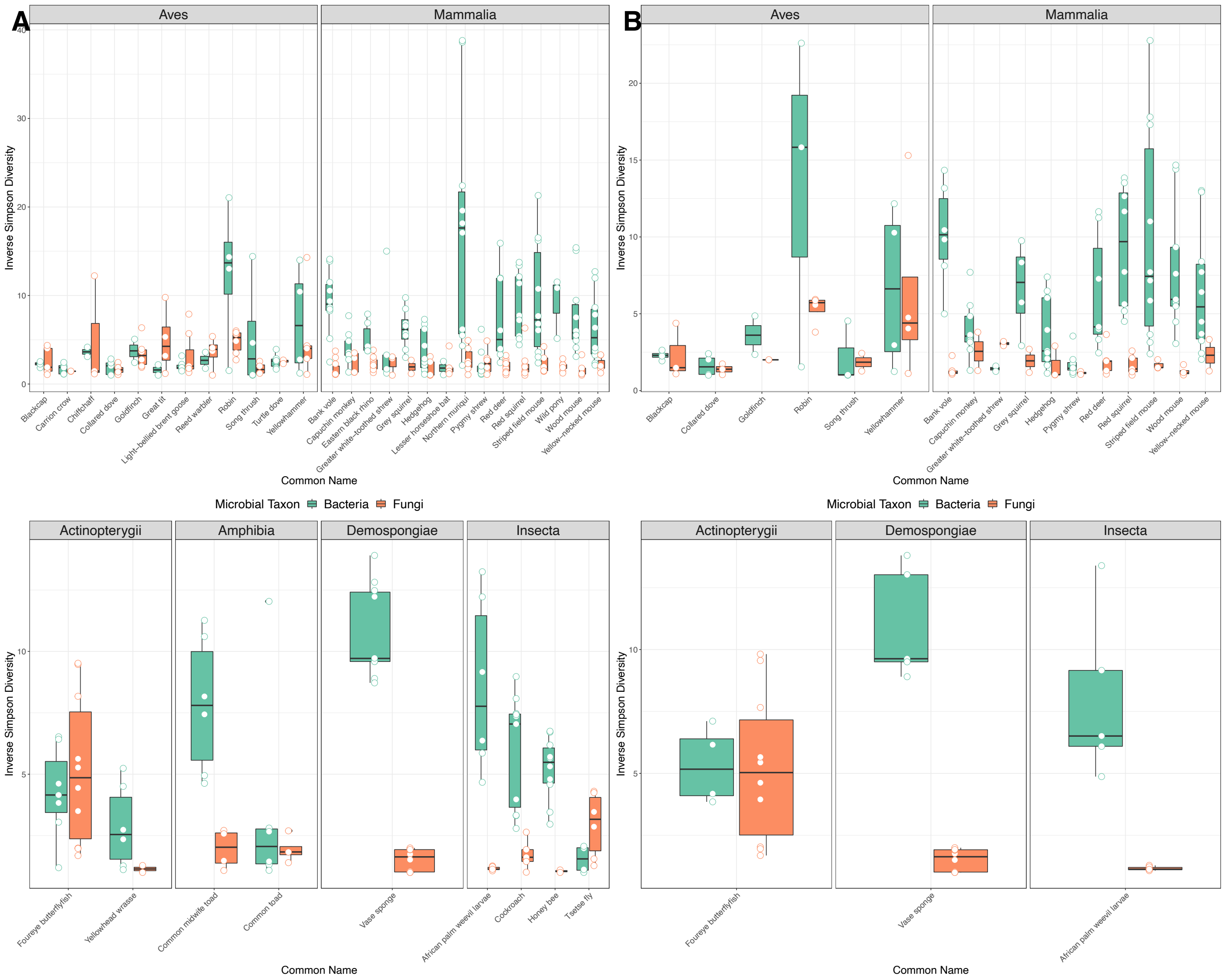


**FIGURE S1: (A)** Median (± $25^{th}$ and $75^{th}$ percentiles) inverse Simpson indices for bacterial (red) and fungal (blue) communities across a range of host species, with amplicon sequence data rarefied to 1000 reads. **(B)** Median (± $25^{th}$ and $75^{th}$ percentiles) inverse Simpson indices for bacterial (red) and fungal (blue) communities across a range of host species, with amplicon sequence data rarefied to 2500 reads. Only species with both bacterial and fungal data are shown.

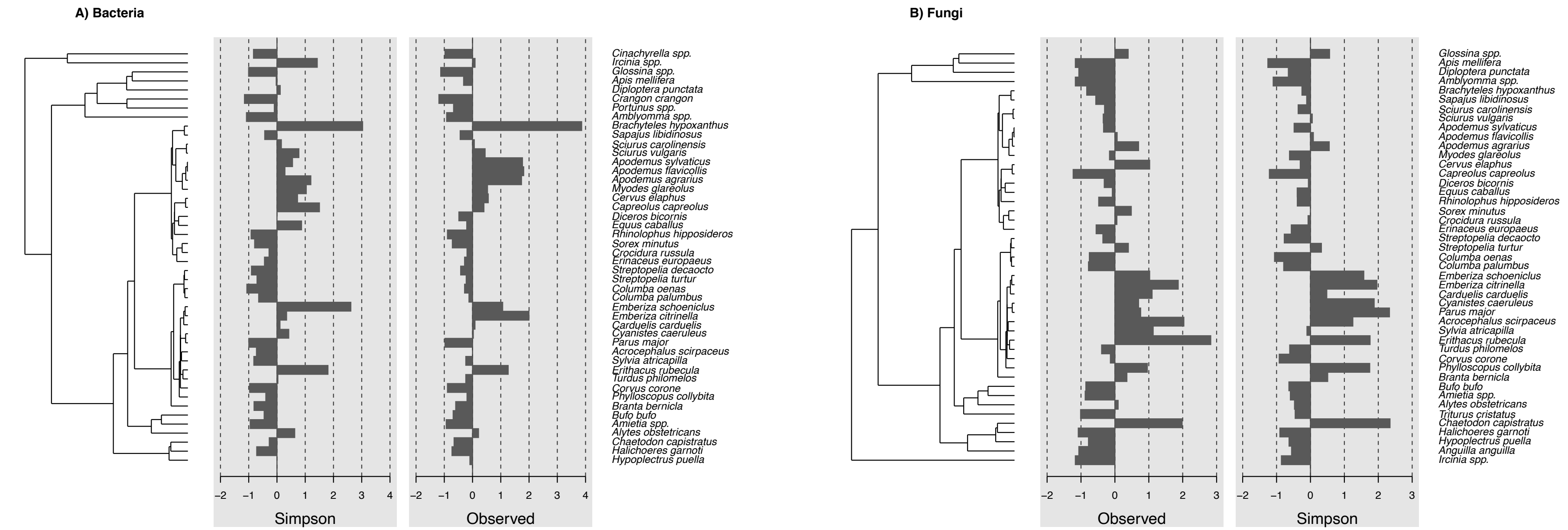


**FIGURE S2.** Patterns of phylogenetic signal in alpha diversity for **(A)** bacteria and **(B)** fungi. Replication of species differs across microbial datasets, reflected by differing host phylogenies.

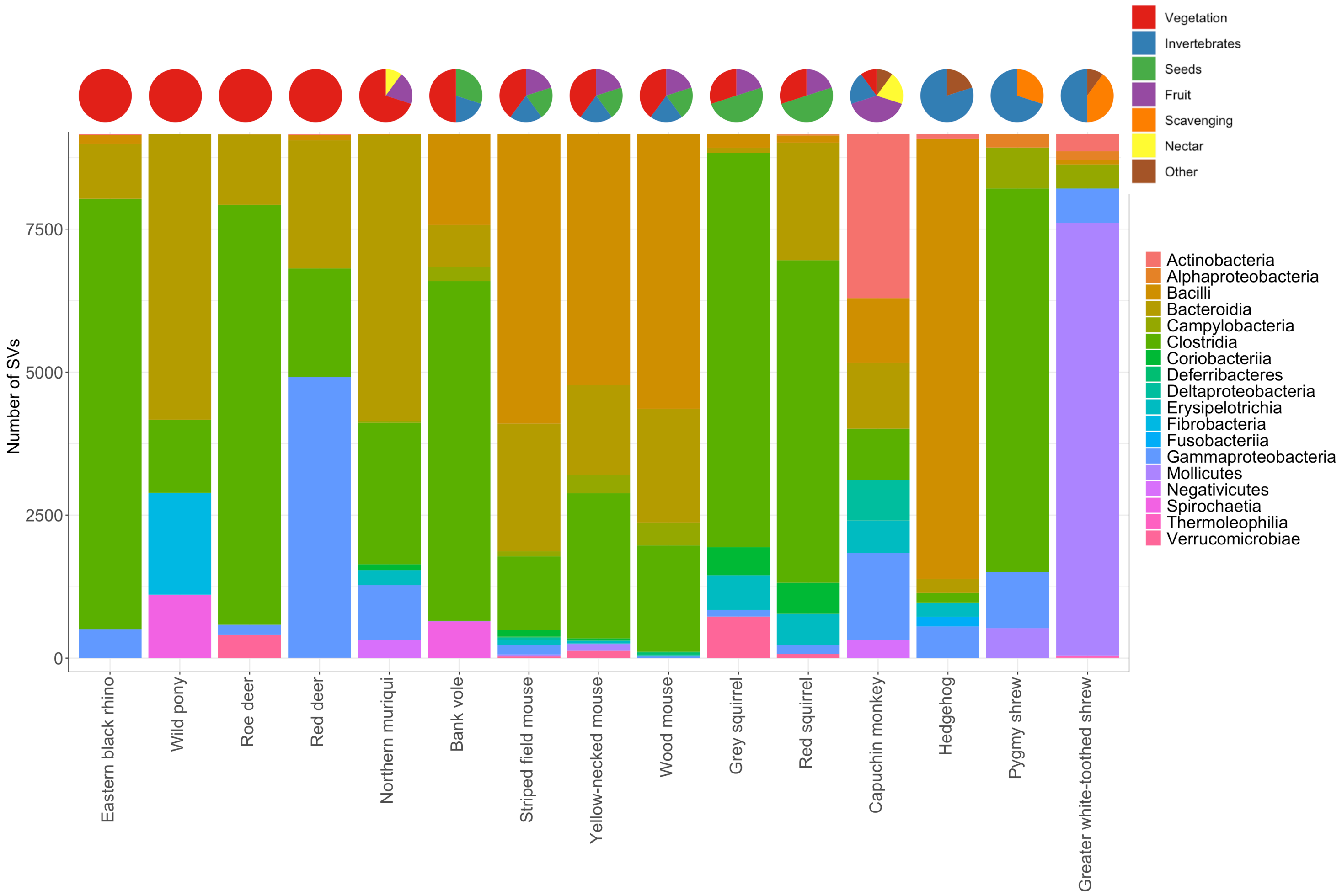


**FIGURE S3:** Bacterial community composition (agglomerated to class level) of 15 mammal species compared with crude foraging data for each host species (see main text for methods and sources). The five most abundant classes of fungi across all host species were Dothideomycetes, Eurotiomycetes, Lecanoromycetes, Pezizomycetes and Sordariomycetes, for which Dothideomycetes and Eurotiomycetes showed the most variation between host species (Fig. S3). The five most abundant classes of bacteria were Actinobacteria, Alphaproteobacteria, Bacilli, Bacteroidia, and Gammaproteobacteria, which all varied considerably among host species.

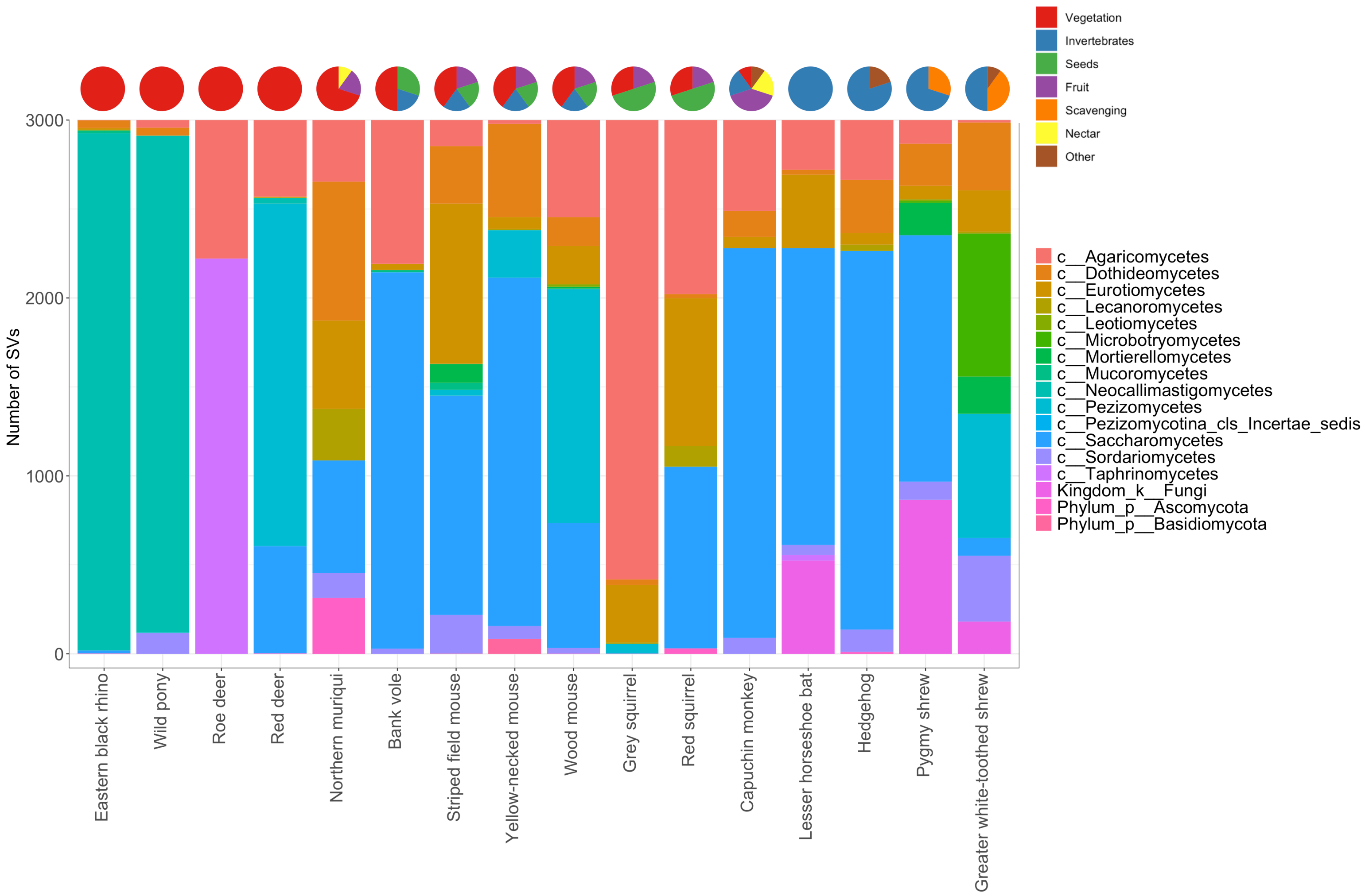


**FIGURE S4:** Fungal community composition (agglomerated to class level) of 16 mammal species compared with crude foraging data for each host species (see main text for methods and sources).

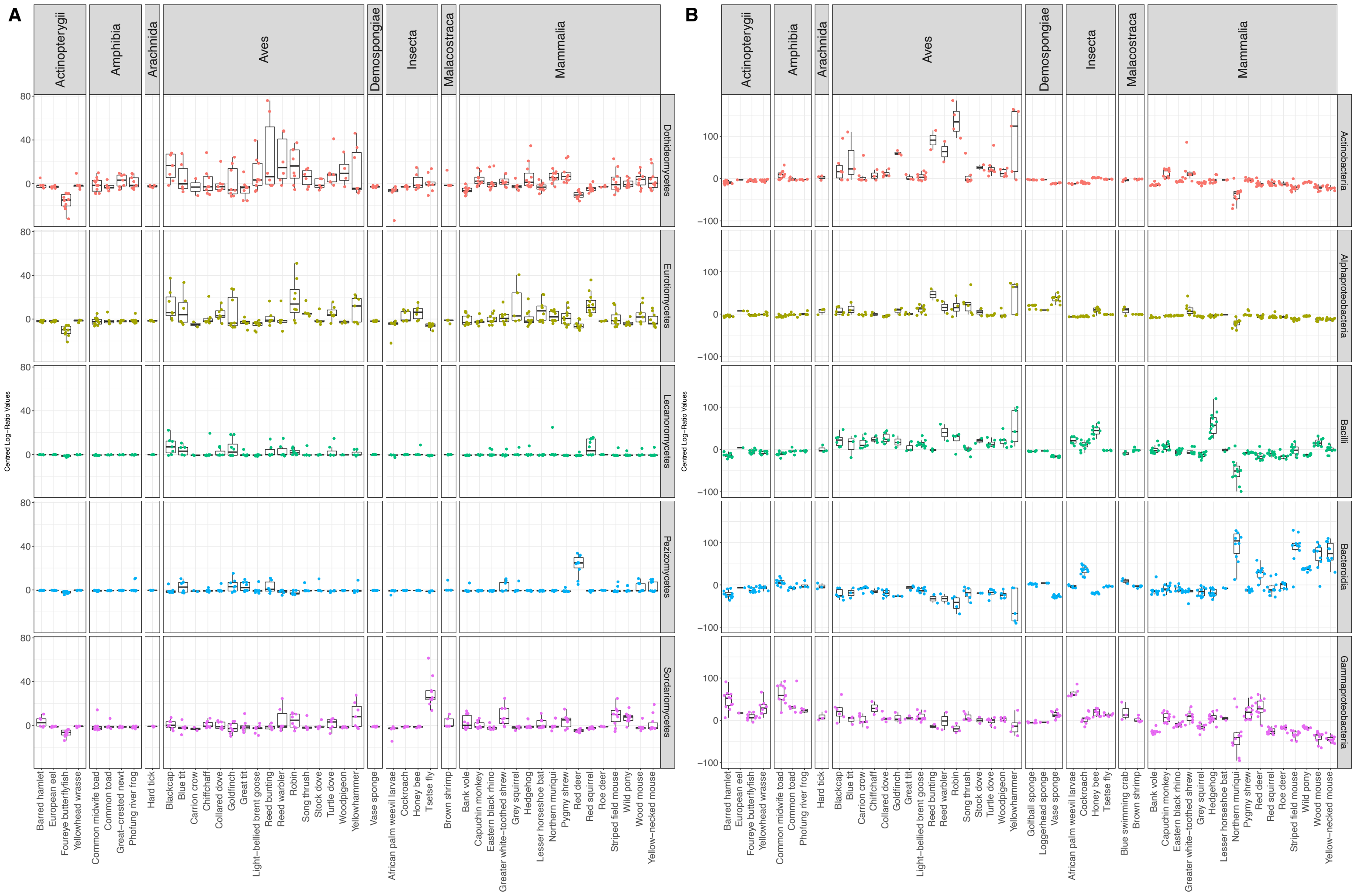


**FIGURE S5.** Centred Log Ratio (CLR)-transformed abundance values from the five most abundant classes of (a) fungi and (b) bacteria identified across a range of host species. CLR-transformation is a normalisation method allowing comparison of abundance values across libraries of different sizes (read depths).

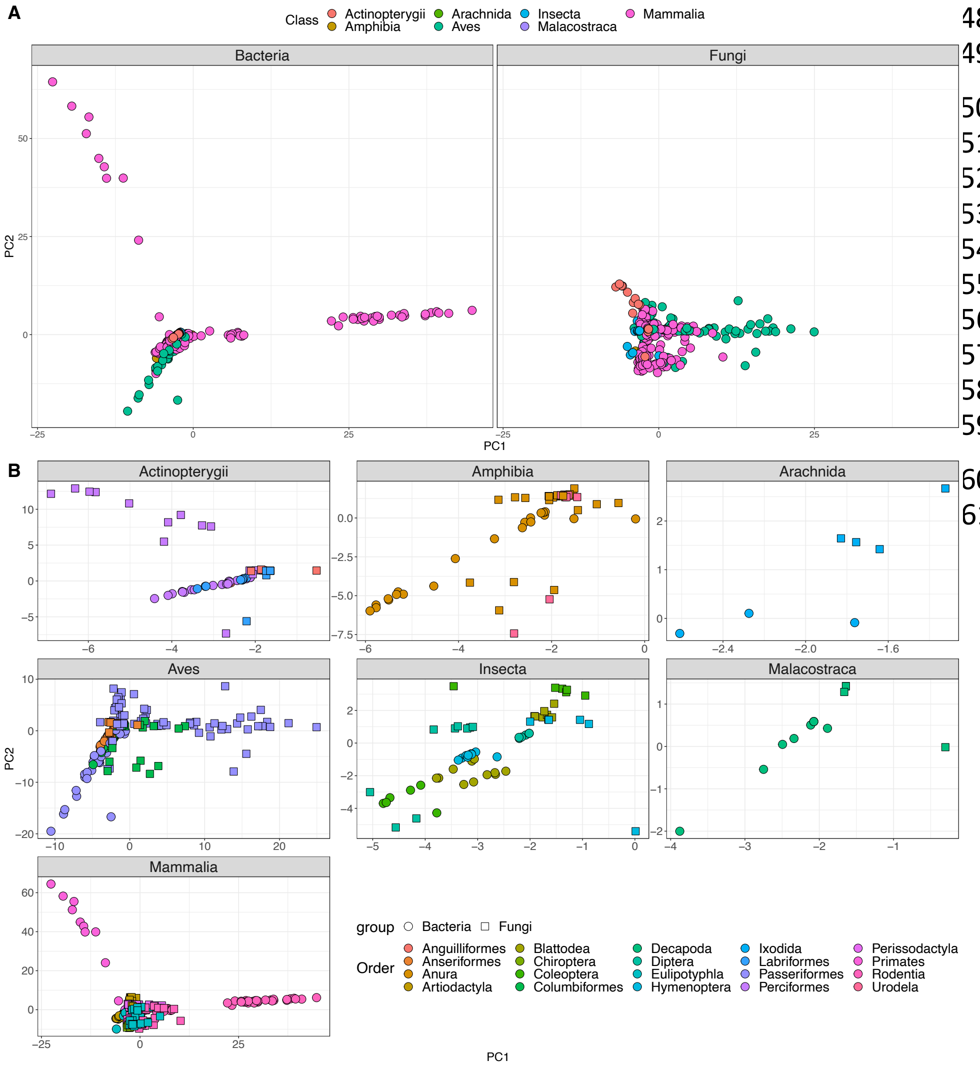


**FIGURE S6**

**(A)** Principal components analysis (PCA) of CLR-transformed microbial abundances for bacteria and fungi, with points coloured by host class. The first two axes of the ordination explained 19.4% and 8.84% of the variance in community structure for bacteria and fungi, respectively. PERMANOVA analysis revealed species ID to be the primary driver of variance in both taxa, accounting for 21.2% and 14.3% of the variance, respectively. However, there were also strong effects of sample handling and storage (see results). **(B)** PCA plots of bacterial (circles) and fungal (squares) community structure, faceted by host class, with points coloured by host order.

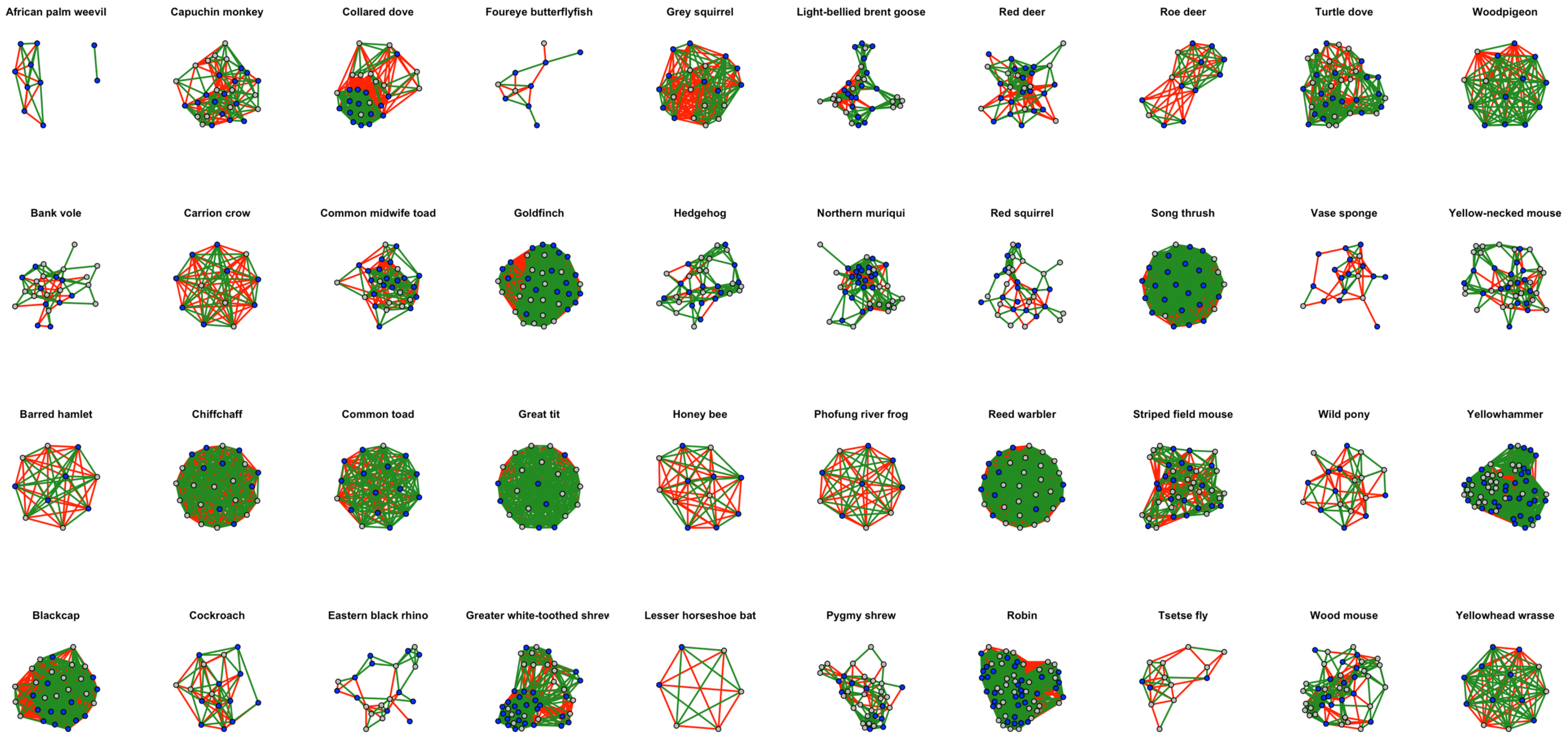


**FIGURE S7:** Microbial interaction networks for 40 species derived from cooccurrence analysis. Positive interactions (correlations) are shown in green, and negative interactions in red; blue nodes are bacteria and grey nodes and fungi. There was clear variation at the species level; for some host species, there were considerably more positive interactions (e.g., yellowhammers, pygmy shrews, greater white-toothed shrews, wood mouse, woodpigeon, yellow-necked mouse. In some species, there were slightly more negative interactions than positive (e.g., blackcap, goldfinch).

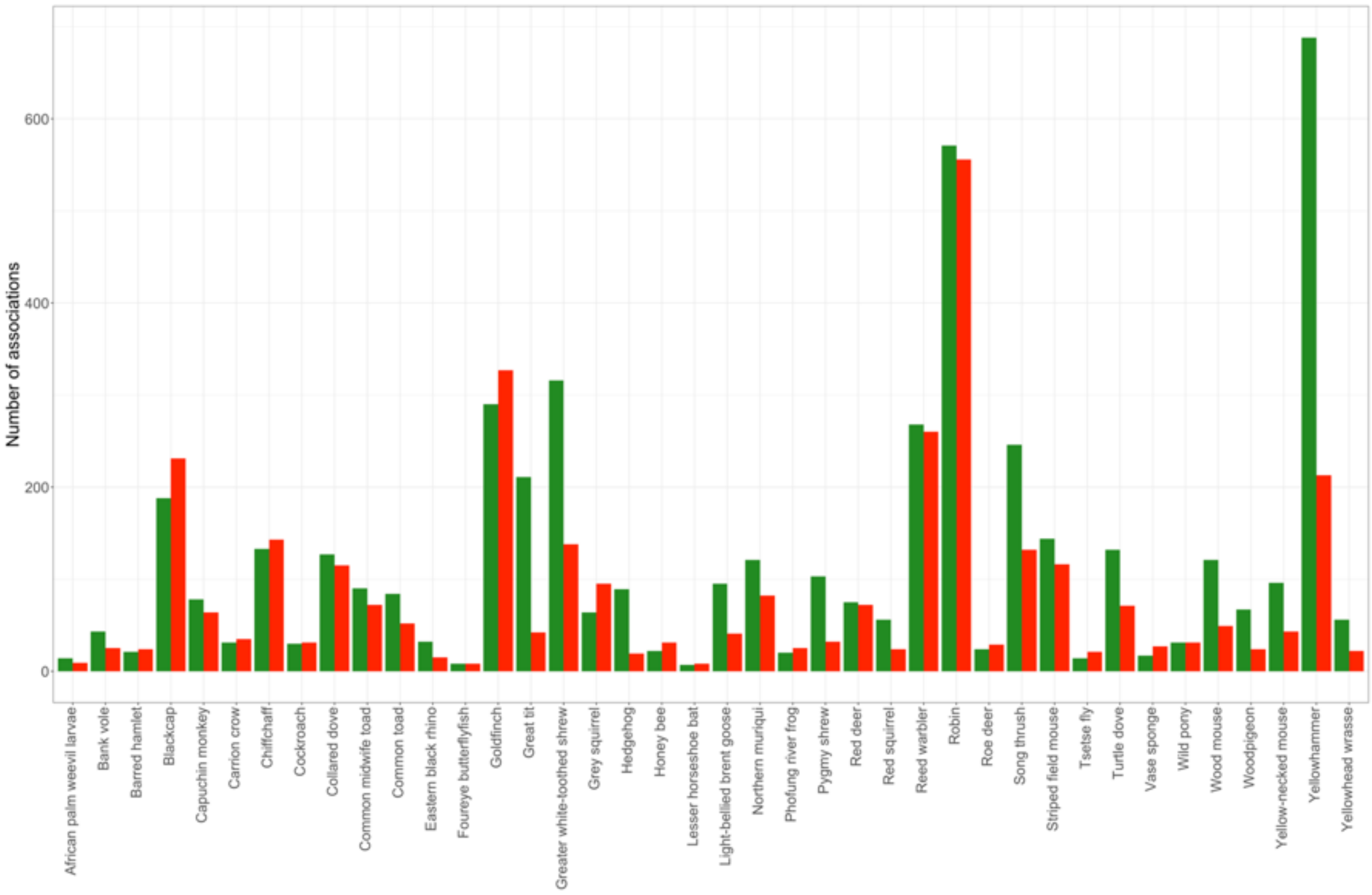


**FIGURE S8:** Number of positive (green) and negative (red) associations within bacterial-fungal co-occurrence networks for each host species.
